# Parallel evolution of frog antimicrobial peptides produces identical conformations but subtly distinct membrane and antibacterial activities

**DOI:** 10.1101/388967

**Authors:** Giorgia Manzo, Philip M. Ferguson, Charlotte Hind, Melanie Clifford, V. Benjamin Gustilo, Hind Ali, Sukhvinder S. Bansal, Tam T. Bui, Alex F. Drake, R. Andrew Atkinson, Mark J. Sutton, Christian D. Lorenz, David A. Phoenix, A. James Mason

## Abstract

Frogs such as *Rana temporaria* and *Litoria aurea* secrete numerous closely related antimicrobial peptides (AMPs) as an effective chemical dermal defence. Despite the high similarity in physical properties and preference for adopting secondary amphipathic, α-helix conformations in membrane mimicking milieu, their spectrum of activity and potency often varies considerably. Damage or penetration of the bacterial plasma membrane is considered essential for AMP activity and hence distinguishing apparently similar AMPs according to their behaviour in, and effects on, model membranes will inform understanding of species specific effective antimicrobial mechanisms. Here we use a combination of molecular dynamics simulations, circular dichroism and patch-clamp to investigate the basis for differing anti-bacterial activities in representative AMPs from each species; temporin L and aurein 2.5. Despite adopting near identical, α-helix conformations in the steady-state in a variety of membrane models, these two AMPs can be distinguished both *in vitro* and *in silico* based on their dynamic interactions with model membranes; the greater conformational flexibility and the higher amplitude channel conductance induced offers a rationale for the greater potency and broader spectrum of activity of temporin L over aurein 2.5. Specific contributions from individual residues are identified that define the mechanisms of action of each AMP. Our findings suggest AMPs in frogs are examples of parallel evolution whose utility is based on apparently similar but subtly distinct mechanisms of action.

## Introduction

Antimicrobial peptides (AMPs) have remained an effective component of the innate immune system throughout evolutionary history and have been isolated from numerous and diverse organisms.^1–6^ Naturally occurring AMPs have attracted considerable interest as a starting point for rational design processes aiming to enhance antimicrobial or immunomodulatory properties.^3–6^ Such AMPs are often found to be part of a larger family of very close-related peptides which share substantial primary sequence and physico-chemical similarities and seemingly have very similar functions. The benefits of producing such a range of similar AMPs remains poorly understood and it is not yet clear the extent to which antimicrobial functions are duplicated by such AMPs. Further, since many AMPs, that share the same physico-chemical properties and have the same antimicrobial function, have been described in unrelated organisms, it is tempting to consider each class of AMP as an example of parallel evolution. Of fundamental interest, therefore, is to what extent such related AMPs can be considered alike or, alternatively, in what ways can their activities and functions be distinguished and explained at a molecular level. In particular, since the ability of AMPs to damage or cross the bacterial plasma membrane is considered to be a major determinant of antibacterial outcomes,^7^ the mechanism by which AMPs interact with models of such membranes is expected to be particularly informative.

Recently we have considered how relatively minor modifications to the primary sequence of temporin B, an AMP from *Rana temporaria*, can fundamentally alter, not only its potency and spectrum of antibacterial activity, but also its interaction and mode of disruption of membranes designed to mimic the plasma membrane of Gram-positive or Gram-negative bacteria.^9^ Interestingly, this study showed that AMPs can be produced that share substantial sequence similarity and that have very similar antibacterial potencies but may nevertheless act with significantly different mechanisms of action. Notably, while biophysical studies conducted in the steady-state struggled to identify any difference in the conformation or membrane interaction of the three temporin B analogues that could be associated with differences in antibacterial activity, time-resolved techniques – notably all-atom MD simulations and patch-clamp – could effectively distinguish the behaviour of the three analogues and provide a rationale for the impact of sequence modification on membrane and antibacterial activities. Previously we used a similar approach to study and compare magainin 2 and pleurocidin. These AMPs have dissimilar primary sequences but share numerous physico-chemical properties, including a preference for α-helix conformation. Nevertheless, substantial differences in the ability of these two AMPs to penetrate model membranes were shown.^10^ This was associated with a greater conformational flexibility of pleurocidin which is consistent with its ability to access intracellular, bactericidal targets and greater potency.^11^

Temporin L is a further example of the ten temporin peptides originally isolated from the European red frog *Rana temporaria*^8^ and has received considerable attention^12^. It is noticeably more potent than other temporin peptides against both Gram-positive and Gram-negative bacteria, including temporin B, but can act in synergy with at least some of these.^13,14^ Its preference for adopting a highly stable α-helix conformation has been described^15^ – again more ordered than temporin B^14^ - and numerous structure-activity-relationship studies have been performed to enhance its antimicrobial^16–20^ or anti-endotoxin^21–24^ activities. AMPs have similarly been isolated from Australian frogs. Seventeen aurein peptides were found in secretions from the granular dorsal glands of the Green and Golden Bell Frog *Litoria aurea*.^25^ Of these, aurein 2.5 has been studied in some detail.^26–30^ Aurein 2.5 also has a strong preference for adopting ordered α-helix conformations in a variety of membrane environments^27,29,30^ and strong surface activity was found in both anionic and zwitterionic model membranes.^26^

*Rana temporaria* and *Litoria aurea* are examples of frogs from, respectively, the Ranidae and Hylidae families. Divergence from their common ancestor occurred during the Jurassic period, approximately 150 million years ago and,^31^ although temporin L and aurein 2.5 are both AMPs derived from frog dorsal secretions, triggered by mild electrical stimulation,^8,25^ they share no sequence similarity. Despite this they do share numerous physico-chemical properties (Table 1); they are both relatively short and hydrophobic, they carry a modest positive charge and are known to adopt ordered α-helix conformations in model membranes or membrane mimicking milieu due to their secondary amphipathicity.^12,27^ Here we consider the antibacterial activity of temporin L and aurein 2.5 and investigate, using an approach that combines both time-resolved and steady-state biophysical methods, whether their membrane interactions can be distinguished. This enables the similarities and some key differences between the two AMPs to be described at the molecular level, allowing better appreciation of the extent to which temporin L and aurein 2.5 may be considered examples of parallel evolution as well as the importance of specific molecular features – notably conformational flexibility – for potent antibacterial activity.

**Table 1.** Peptides sequences, biophysical characteristics and concentration of peptide necessary to start membrane activity in electrophysiology experiments. Both peptides were amidated at the C-terminus. Hydrophobicity (H) and hydrophobic moment (µH) were calculated using HeliQuest.82

| Peptide | Sequence | Length | Charge | H | $\mu H$ | Peptide concentration ($\mu M$) | |
| --- | --- | --- | --- | --- | --- | --- | --- |
|  |  |  |  |  |  | DPhPE/DPhPG | DPhPG |
| Aurein 2.5 | GLF**D**IV**KK**VV GAFGSL | 16 | +2 | 0.622 | 0.609 | 7.69 | 5.75 |
| Temporin L | FVQWFS**K**FLG **R**IL | 13 | +3 | 0.906 | 0.710 | 9.52 | 10 |

## Experimental procedures

### Peptides and lipids

Peptides (Table 1) were purchased from Pepceuticals Ltd (Enderby, UK) or Cambridge Research Biochemicals (Cleveland, UK) as desalted grade (crude) and were further purified using water/acetonitrile gradients using a Waters SymmetryPrep C8, 7 mm, 19 x 300 mm column. They were both amidated at the C-terminus. The lipids 1-palmitoyl-2-oleoyl-*sn*-glycero-3-phospho-(1′-*rac*-glycerol) (POPG), 1-palmitoyld31-2-oleoyl-*sn*-glycero-3-phospho-(1′-*rac*-glycerol) (POPG-d31), 1-palmitoyl-2-oleoyl-*sn*-glycero-3-phosphoethanolamine (POPE), 1-palmitoyld31-2-oleoyl-*sn*-glycero-3-phosphoethanolamine (POPE-d31), 1,2-diphytanoyl-*sn*-glycero-3-phospho-(1'-*rac*-glycerol) (DPhPG) and 1,2-diphytanoyl-*sn*-glycero-3-phosphoethanolamine (DPhPE) were purchased from Avanti Polar Lipids, Inc. (Alabaster, AL) and used without any purification. All other reagents were used as analytical grade or better.

### Antibacterial activity assay

The antibacterial activity of the five peptides was assessed through a modified two-fold broth microdilution assay, as described before^10^. Briefly, a two-fold dilution of peptide stock solutions was performed in non-cation adjusted Mueller Hinton broth in a 96-well polypropylene microtiter plate. Then, 100 µl of bacterial suspension (back diluted from overnight cultures to an A_600_ of 0.01) were added to 100 µl of peptide solution in each well of the plate. Growth and sterility controls were present for each experiment. The plate was incubated at 37°C for 18 hours without shaking and the MIC was defined as the lowest concentration where no visible growth could be detected by measuring the absorbance (A_600_) on an Omega plate reader.

### NMR structure determination

Liquid state NMR spectroscopy experiments and data analysis were performed as described previously^10^. The samples solution consisted of 2 mM peptide and 100 mM deuterated sodium dodecyl sulphate (SDS-d25) micelles. 10% D_2_O containing trimethylsilyl propanoic acid (TSP) was added for the lock signal and as internal chemical shift reference. NMR spectra were acquired on a Bruker Avance 500 MHz spectrometer (Bruker, Coventry, UK) equipped with a cryoprobe. Standard Bruker TOCSY and NOESY pulse sequences were used, with water suppression using an excitation sculping sequence with gradients (mlevesgpph and noesyesgpph). The ^1^H 90-degree pulse length was 8.0 s. One TOCSY with a mixing time of 60 ms, and two NOESY with mixing time of 100 and 200 ms were acquired for each sample. The relaxation delay was 1 s. 2048 data points were recorded in the direct dimension, and either 256 or 512 data points in the indirect dimension. CARA (version 1.9.1.2) and Dynamo software^30,31^ were used for the structure calculation. Inter-proton NOEs interactions were used as distance restraints in the structure calculation. CARA software generated a total of 200 structures on UNIO’08 (version 1.0.4)^34^ and XPLOR-NIH (version 2.40)^35,36^, after 7 iterations, using the simulated annealing protocol. 20 structures with the lowest energy were chosen to produce a final average structure. In the case of ambiguous NOEs assignments by CARA software, Dynamo software’s annealing protocol was applied. Only unambiguous NOEs were used in this case after being classified as strong, medium and weak on the base of the relative intensity of the cross-peaks in the NOESY spectra. On the base of this classification an upper limit of 0.27, 0.33 and 0.50 nm have been applied, respectively, as restraint on the corresponding inter-proton distance, as described previously^37^. One thousand structures were calculated and the 100 conformers with the lowest potential energy were selected for the analysis. The selected 100 conformers were aligned, and the root mean square deviation (RMSD) of the backbone heavy atoms was calculate with respect to their average structure. Solvent molecules were not included in the calculations. Structural coordinates were deposited in the Protein Data Bank (www.rcsb.org) under accession codes 6GS5 and 6GS9 for temporin L and aurein 2.5 respectively.

### Molecular dynamics simulations

Simulations were carried out on either a Dell Precision quad core T3400 or T3500 workstation with a 1 kW Power supply (PSU) and two NVIDA PNY GeForce GTX570 or GTX580 graphics cards using Gromacs^38^. The CHARMM36 all-atom force field was used in all simulations^39,40^. The initial bilayer configuration was built using CHARMM-GUI^41^. All membranes in this project contained a total of 512 lipids, composed either of POPE/POPG (75:25 mol:mol) or POPG to reflect the lipid charge ratios of the plasma membrane of Gram-negative and Gram-positive bacteria, respectively. Eight peptides were inserted at least 30 Angstrom above the lipid bilayer in a random position and orientation at least 20 Angstrom apart. The starting structures were taken from the NMR calculation in SDS micelles. The system was solvated with TIP3P water and Na+ ions added to neutralize. Energy minimization was carried out at 310 K with the Nose-Hoover thermostat using the steepest descent algorithm until the maximum force was less than 1000.0 kJ/ml/nm (~3000-4000 steps). Equilibration was carried out using the NVT ensemble for 100 ps and then the NPT ensemble for 1000 ps with position restraints on the peptides. Hydrogen-containing bond angles were constrained with the LINCS algorithm. Final simulations were run in the NPT ensemble using 2 femtosecond intervals, with trajectories recorded every 2 picoseconds. All simulations were run for a total of 100 nanoseconds and repeated twice, with peptides inserted at different positions and orientations, giving a total of approximately 0.8 µs simulation.

### Liposome preparation

Small unilamellar vesicles (SUVs) and multi-lamellar vesicles (MLVs) were prepared for circular dichroism (CD) as previously described^10^. Lipid powders were solubilized in chloroform and dried under rotor-evaporation. To completely remove the organic solvent, the lipid films were left overnight under vacuum and hydrate in 5 mM Tris buffer with or without the addition of 100 mM NaCl (pH 7.0). Lipid suspension was subjected to 5 rapid freeze-thaw cycles for further sample homogenisation. POPE/POPG (75:25, mol:mol) and POPG SUVs were obtained by sonicating the lipid suspension on Soniprep 150 (Measuring and Scientific Equipment, London, UK) for 3 × 7 minutes with amplitude of 6 microns in the presence of ice to avoid lipid degradation. The SUVs were stored at 4C_∘_ and used within 5 days of preparation.

### Circular dichroism spectroscopy

Far-UV spectra of the peptides in the presence of SUVs and SDS micelles were acquired on a Chirascan Plus spectrometer (Applied Photophysics, Leatherhead, UK). Liposome samples were maintained at 310 K. Spectra were recorded from 260 to 190 nm. Lipid suspension was added to a 0.5 mm cuvette at a final concentration of 5.0 mM and then a few μl of a concentrated peptide solution were added and thoroughly mixed to give the indicated final peptide-to-lipid molar ratios. The same experimental conditions were used to investigate peptides secondary structure in SDS micelles. Final peptide concentration in the 0.5 mm cuvette was 40 μM, while SDS micelles concentration was 2 mM (L/P=50). In processing, a spectrum of the peptide free suspension was subtracted and Savitsky-Golay smoothing with a convolution width of 5 points applied.

### Electrophysiology experiments (Patch-clamp)

Giant unilamellar vesicles (GUVs) composed of DPhPE/DPhPG (60:40, mol:mol) and DPhPG were prepared in the presence of 1 M sorbitol by the electroformation method in an indium-tin oxide (ITO) coated glass chamber connected to the Nanion Vesicle Prep Pro setup (Nanion Technologies GmbH, Munich, Germany) using a 3-V peak-to-peak AC voltage at a frequency of 5 Hz for 120 and 140 minutes, respectively, at 36°C^42–44^. Bilayers were formed by adding the GUVs solution to a buffer containing 250 mM KCl, 50 mM MgCl_2_ and 10 mM Hepes (pH 7.00) onto an aperture in a borosilicate chip (Port-a-Patch®; Nanion Technologies) and applying 70-90 mbar negative pressure resulting in a solvent-free membrane with a resistance in the GΩ range. After formation, a small amount of peptide stock solution (in water) was added to 50 μL of buffer solution in order to obtain its active concentration. All the experiments were carried on with a positive holding potential of 50 mV. The active concentration, the concentration at which the peptide first showed membrane activity, for each peptide was obtained through a titration performed in the same conditions. For all the experiments a minimum of 6 according repeats was done. Current traces were recorded at a sampling rate of 50 kHz using an EPC-10 amplifier from HEKA Elektronik (Lambrecht, Germany). The system was computer controlled by the PatchControl™ software (Nanion) and GePulse (Michael Pusch, Genoa, Italy, http://www.ge.cnr.it/ICB/conti_moran_pusch/programs-pusch/software-mik.htm). The data were filtered using the built-in Bessel filter of the EPC-10 at a cut-off frequency of 10 kHz. The experiments were performed at room temperature. Data analysis was performed with the pClamp 10 software package (Axon Instruments). Estimation of pore radii was performed as previously.^45^

## Results

### Temporin L is a more potent antimicrobial than aurein 2.5

In the present study, temporin L is shown to be more potent than aurein 2.5 against all strains included in both the Gram-positive and Gram-negative bacteria panels and *Candida albicans* (Table 2). For the Grampositive strains, aurein 2.5 is outperformed by temporin L with, on average a 7.6-fold greater potency for the latter. For the Gram-negative strains, the difference between the potency of aurein 2.5 and temporin L is less, notably for both *Acinetobacter baumannii* isolates. However, the spectrum of activity for aurein 2.5 is narrower with almost no detectable activity against either *Pseudomonas aeruginosa* isolate. If the antibacterial activity of these two AMPs is dependent on their membrane interactions, there is therefore reason to expect differing interactions and behaviours of each AMP with model membranes designed to mimic the differing plasma membranes of these two groups.

**Table 2.** **Antimicrobial activity**. MS – methicillin sensitive; EMR – epidemic methicillin resistant; VS – vancomycin sensitive; VR – vancomycin resistant.

| | Isolate | Peptide concentration ($\mu g/ml$) | |
| --- | --- | --- | --- |
|  |  | Aurein 2.5 | Temporin L |
| Gram-negative | *Klebsiella pneumoniae* NCTC 13368 | 64 | 16 |
|  | *Klebsiella pneumoniae* M6 | 32-64 | 16 |
|  | *Acinetobacter baumannii* AYE | 4-8 | 4 |
|  | *Acinetobacter baumannii* ATCC 17978 | 8 | 4 |
|  | *Pseudomonas aeruginosa* PAO1 | 128 | 16 |
|  | *Pseudomonas aeruginosa* NCTC 13437 | 128 | 32-64 |
|  | *Escherichia coli* NCTC 12923 | 16 | 4 |
| Gram-positive | MS *Staphylococcus aureus* ATCC 9144 | 8 | 2 |
|  | EMR *Staphylococcus aureus*-15 | 32 | 4 |
|  | EMR *Staphylococcus aureus*-16 | 32-64 | 4 |
|  | VS *Enterococcus faecalis* NCTC 775 | 32-64 | 4-8 |
|  | VR *Enterococcus faecalis* NCTC 12201 | 64 | 8 |
| Yeast | *Candida albicans* NCPF 3179 | 16 | 8 |

### Both temporin L and aurein 2.5 adopt highly ordered α-helix conformations in membrane mimicking media

Far-UV circular dichroism spectra were obtained in anionic micelles (Fig. 1C) but also in small lamellar vesicles (SUVs) whose composition was designed to mimic the plasma membrane of Gram-positive or Gram-negative bacteria (Fig. S1). The structure of temporin L has previously been determined both in SDS and dodecylphosphocholine (DPC) micelles^19^ but coordinates are not available in the PDB, while no structure has been determined for aurein 2.5. Structures of both peptides were therefore also solved in the presence of SDS (Fig. 1A/B) and were used subsequently as starting structures for the molecular dynamics simulations. In SDS, both the far-UV CD and the NMR structures agree that both peptides adopt ordered α-helix conformations. The NMR structures and CD spectra can be compared with those obtained for temporin B and its analogues which describe peptides with a much greater degree of disorder.^9^

**Figure 1.**
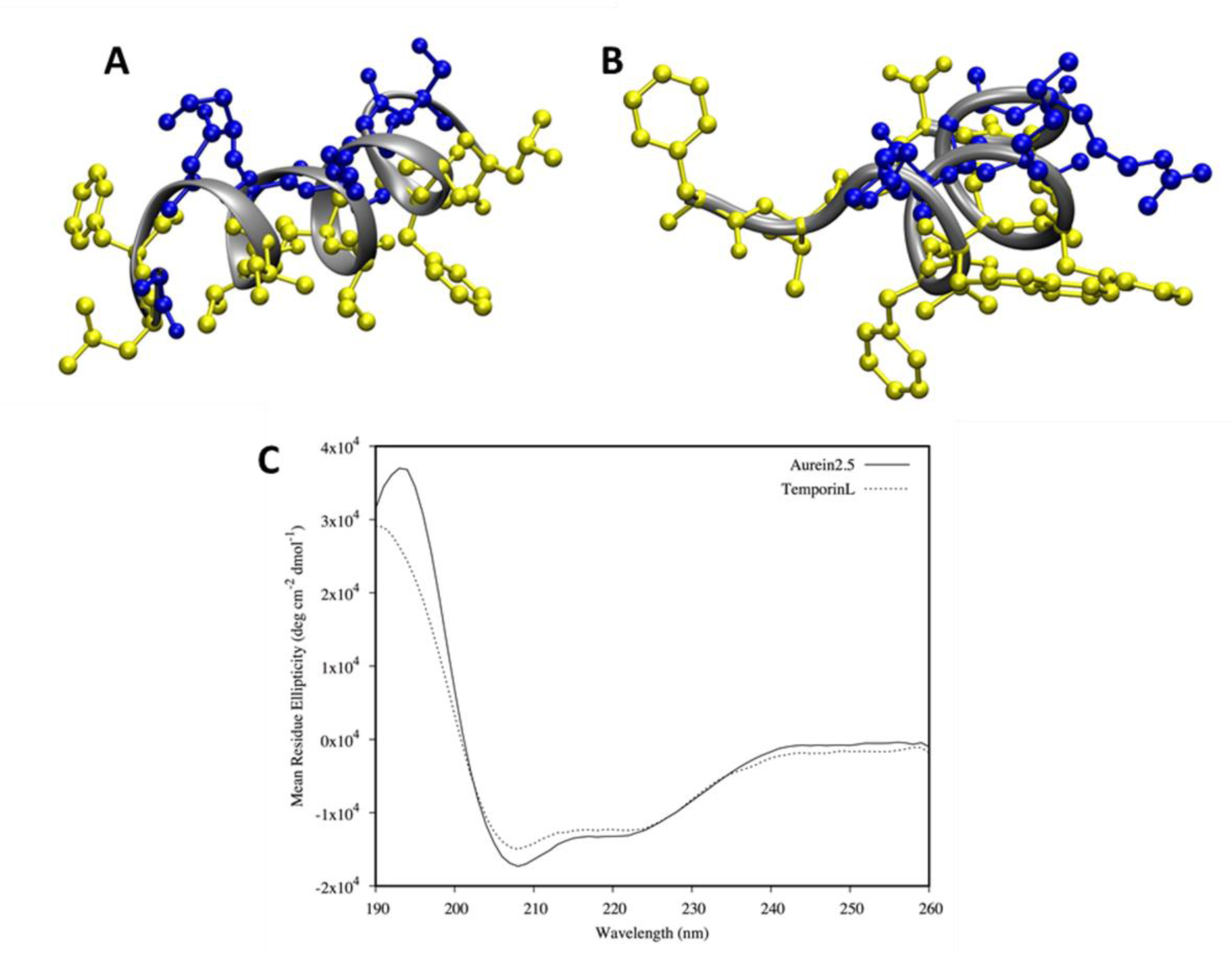
Three-dimensional structure for aurein 2.5 and temporin L. Structures were determined through ^1^H-NOESY NMR spectroscopy in SDS-d25 micelles (peptide/detergent equal to 1/50). The figure shows the structure with the minimum RMSD, used as starting point in the MD simulations. The hydrophobic and hydrophilic residues are shown in yellow and blue for aurein 2.5 (A) and temporin L (B). CD spectra of the peptides in the same conditions used for NMR experiments (C).

The α-helix of aurein 2.5 extends almost throughout the length of the peptide while, for temporin L a little disorder is observable at the N-terminus only. Both peptides display characteristic secondary amphipathicity when adopting this ordered conformation. After binding to lipid bilayers, in the steady-state far-UV CD again indicates similar ordered α-helix conformations are adopted by both peptides (Fig. S1); in the absence of lipid both peptides adopt disordered conformations. There are no apparent differences in conformation between either peptide in SUVs designed to mimic Gram-negative (Fig. S1A/B) or Gram-positive (Fig. S1C/D) plasma membranes and there was no significant effect of varying the peptide to lipid ratio.

### MD simulations reveal conformational flexibility to temporin L but not aurein 2.5 on initial binding to model membranes

Conformational flexibility has proven to be a key parameter capable of distinguishing the behaviour of AMPs with models of both Gram-negative and Grampositive bacteria plasma membranes.^9,10^ Therefore all-atom MD simulations were used to characterise the atomistic details of the interaction and conformational flexibility of aurein 2.5 and temporin L with model bacterial membranes.

Averaged psi dihedral angles for each residue in the eight aurein 2.5 peptides are similar in the presence of both POPE/POPG (Fig. 2A) and POPG (Fig. 2B) membranes; as in previous work,^9,10^ phi dihedral angles were not affected and are not shown. The values for these psi dihedral angles lie in the range expected for α-helix conformation and this is maintained throughout the length of the peptide, even at the N-and C-termini. As evidenced by the relatively low values obtained for the circular variance throughout the length of the peptide, conformational flexibility is modest, and no single region can be identified as being more flexible than another.

**Figure 2.**
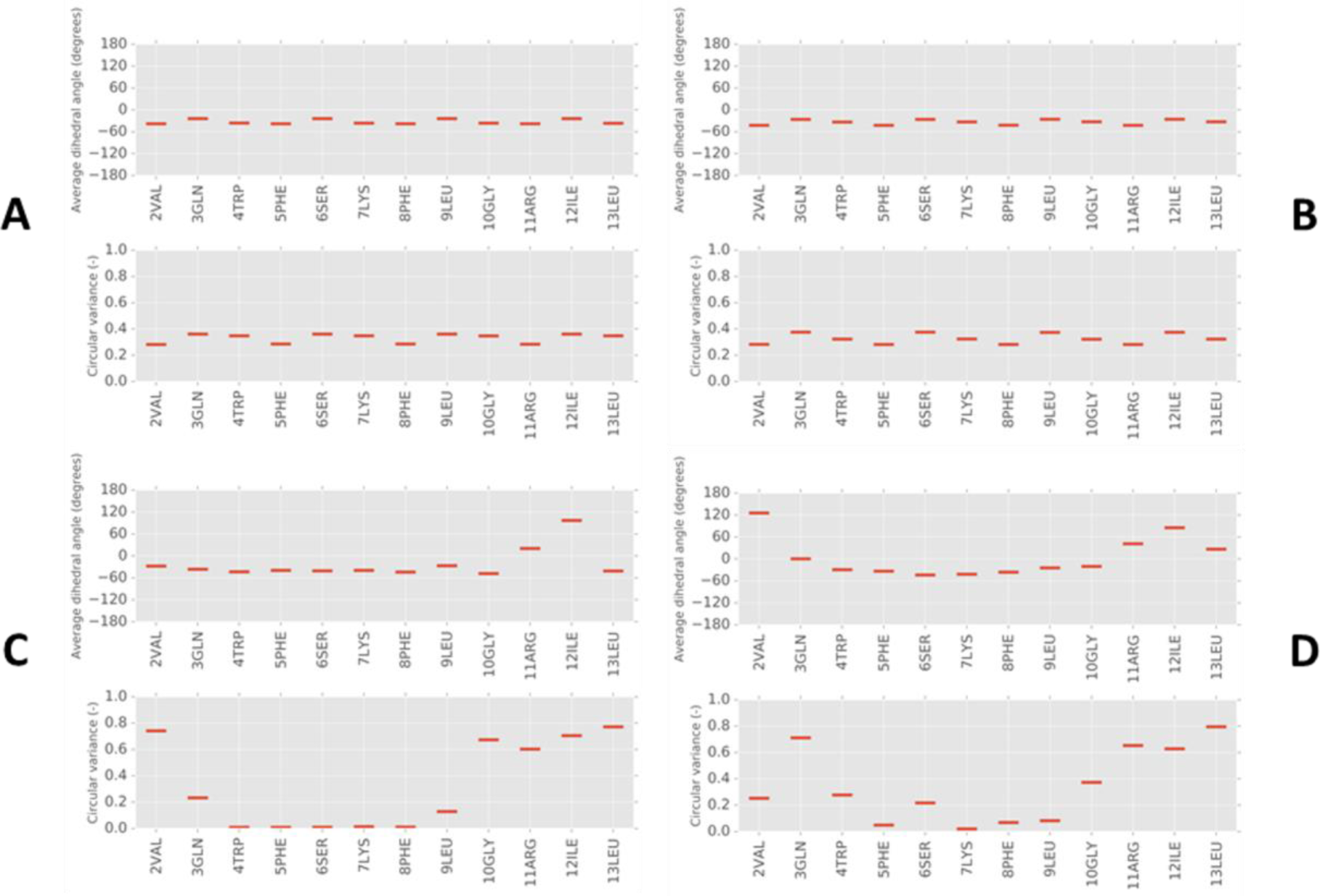
Secondary structure analysis of aurein 2.5 and temporin L peptides from MD simulations. Dihedral angles (psi) and their circular variance are shown for each residue averaged over 100 ns of simulation and eight peptides in aurein 2.5 (A/B) and temporin L (C/D) peptides when binding to POPE/POPG (A/C) or POPG (B/D) membranes. Backbone dihedral angles (psi) for residues in α-helix conformation are around 45°.

In the analogous simulations performed for temporin L however, features are revealed that now distinguish its behaviour in model membranes from aurein 2.5 (Fig. 2C/D). In both POPE/POPG (Fig. 2C) and POPG (Fig. 2D) the α-helix conformation is retained only for the first ten (of thirteen) residues. The three residues that follow Gly10 (Arg11, Ile12 and Leu13) may adopt more extended conformations and the circular variance indicates that the four residues at the C-terminus experience substantial conformational flexibility. Further conformational flexibility is also apparent at the N-terminus, but this does not extend beyond the first two or three residues and, between Trp4 and Leu9, there is a section of highly ordered α-helix (WFSKFL) with almost no variation in psi dihedral angle detected. There is little apparent difference between the secondary structure adopted by temporin L during the initial binding to either POPE/POPG or POPG bilayers apart from perhaps slightly greater conformational flexibility at the N-terminus when binding to POPG (Fig. 2D).

The topology of temporin B during the initial binding to lipid bilayers is sensitive to changes in primary sequence.^9^ Here the ability of aurein 2.5 or temporin L to penetrate POPE/POPG or POPG bilayers can be visualized (Fig. 3; Fig. S2). The two peptides both attempt to penetrate either POPE/POPG or POPG bilayers via the N-terminus. For aurein 2.5 the differing bilayer composition did not appear to have any substantial impact on the ability of the peptide to penetrate and the first six to nine residues were consistently able to penetrate below the plane of the lipid phosphate groups (Fig. 3A/B).

**Figure 3.**
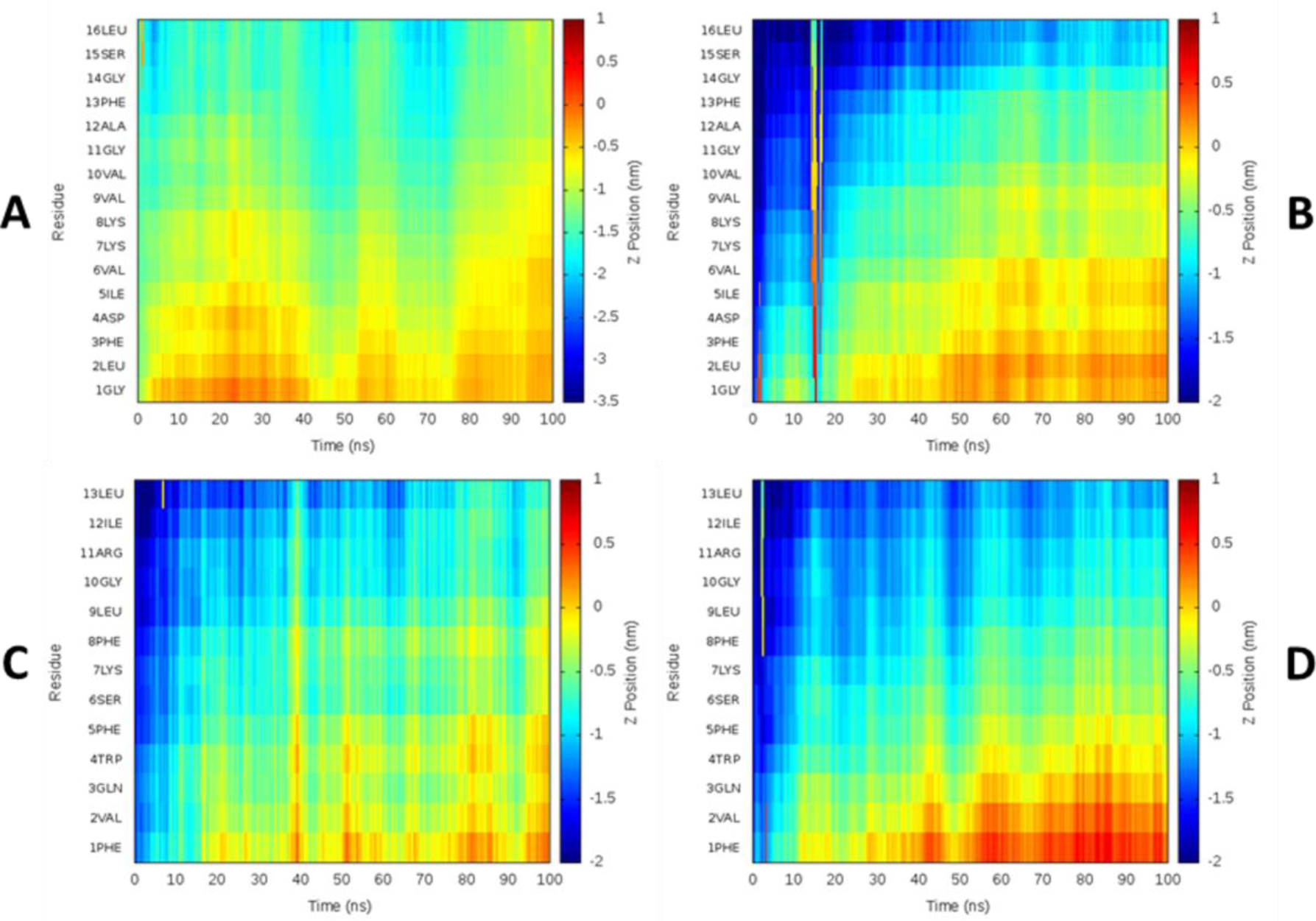
Aurein 2.5 and temporin L peptide topology during 100 ns MD simulation. The depth of aurein 2.5 (A/B) and temporin L (C/D) insertion is shown as the Z-position relative to the phosphate group for each residue averaged over all eight peptides in each simulation. Positive or negative values indicate the peptides are below or above the phosphate group, respectively for peptides in POPE/POPG (A/C) or POPG (B/D).

In contrast, temporin L penetrates POPE/POPG and POPG bilayers in subtly different ways (Fig. 3C/D). In both cases, penetration proceeds via the N-terminus. When binding to POPE/POPG, the first eight residues of temporin L penetrate to the plane of the lipid phosphate and a certain periodicity, consistent with the ordered α-helix conformation, can be observed indicating an alignment roughly parallel to the bilayer surface (Fig. 3C). When binding to POPG however, a deeper penetration by the first four residues at the N-terminus is observed. Resides 5-8 (FSKF) penetrate more slowly and the periodicity is absent, indicating a steeper angle of attack on the bilayer (Fig. 3D). In both cases, the four residues at the N-terminus, which experience conformational flexibility, do not penetrate into the hydrophobic core of the bilayer over the 100 ns of the simulation.

### Hydrophobic amino acids mediate peptide aggregation at the bilayer surface

Both aurein 2.5 (Fig. 4) and temporin L (Fig. 5) were frequently observed to self-associate (Fig. S3). In both POPE/POPG and POPG, aurein 2.5 was observed as a monomer and in dimer or trimer self-associations (Fig. 4B/E). For both dimer (Fig. 4A) and trimer (Fig. 4D) configurations, the self-association was mediated by hydrophobic groups, notably Phe3 and Phe13 which are located near to N-and C-termini respectively and the three valines that are located in the centre of the peptide (Fig. 4C/F). This configuration was qualitatively the same when inserting into both POPE/POPG and POPG membranes and was reproduced in both duplicates.

**Figure 4.**
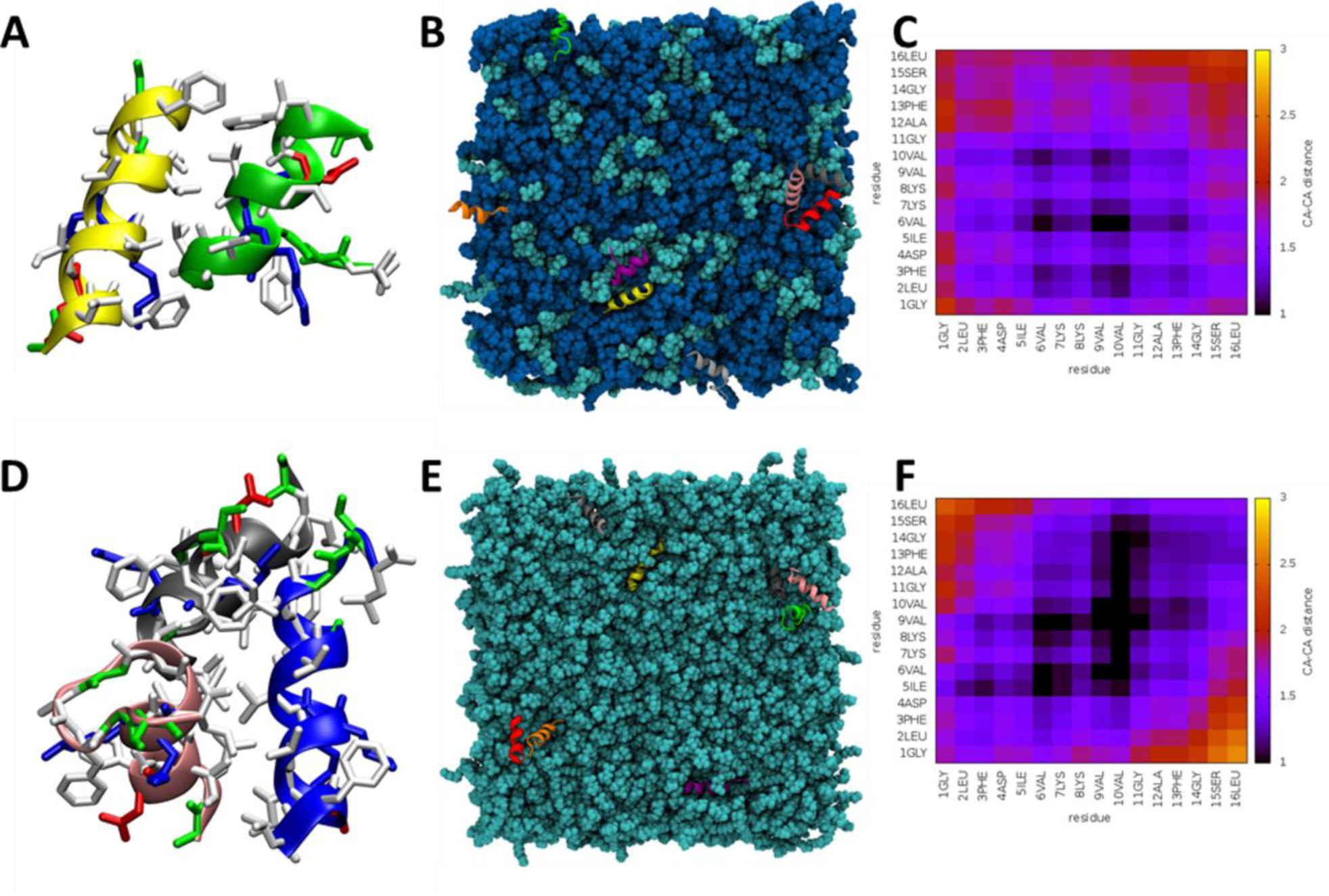
Interaction of aurein 2.5 with lipid bilayers. Examples of peptides aggregating as a dimer or a timer are shown respectively for POPE/POPG (A) and POPG (D). Basic, polar, acidic and hydrophobic residues are coloured in blue, green, red and white respectively. The top view snapshot shows the tendency to aggregate for the eight peptides in POPE/POPG (B) or POPG (E). Heatmaps of average Cα-Cα distances for peptides in POPE/POPG (C) or POPG (F) show the stronger interactions that mediate assembly of aggregates

**Figure 5.**
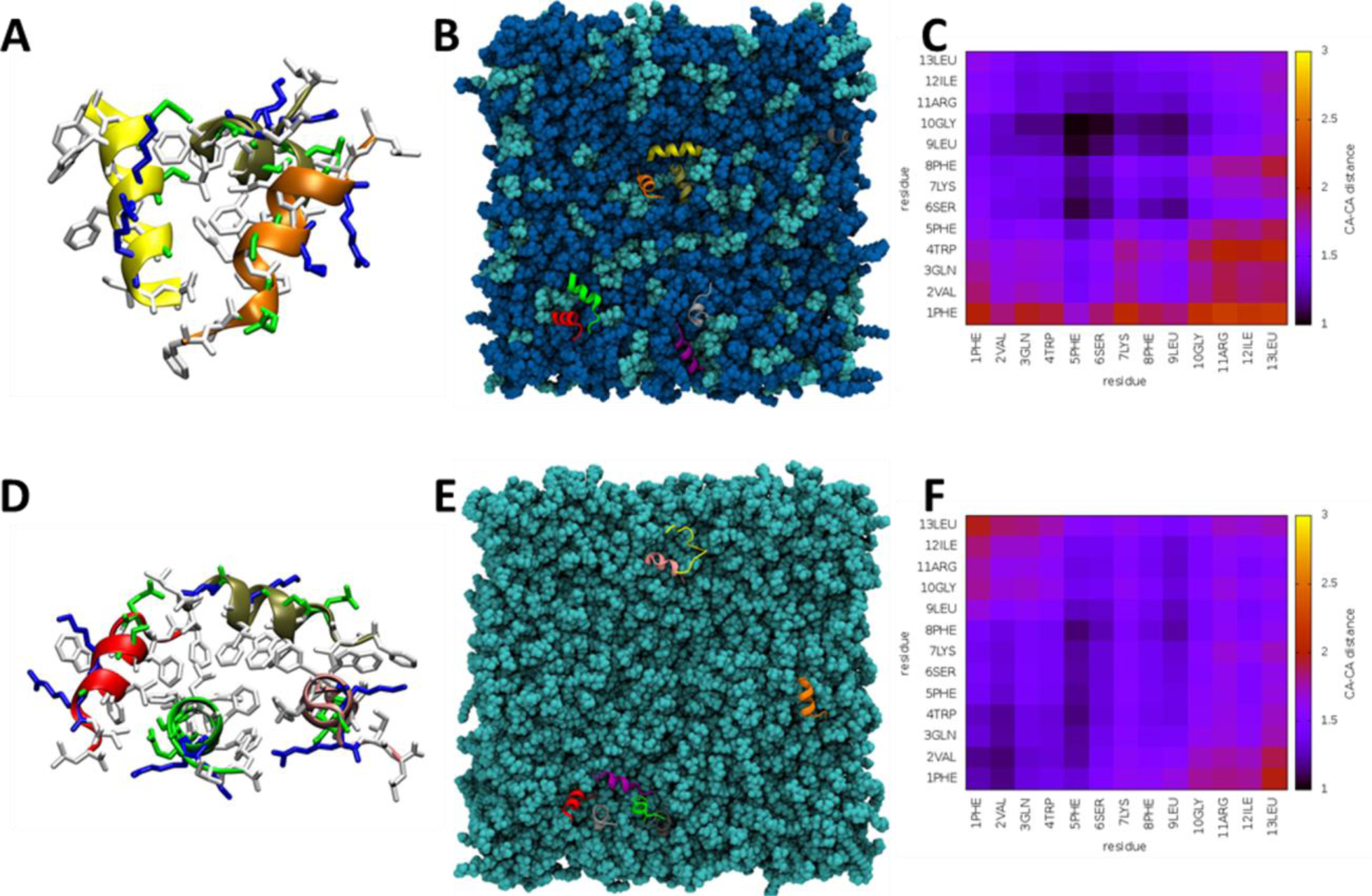
Interaction of temporin L with lipid bilayers. Examples of peptides aggregating as a trimer or a tetramer are shown respectively for POPE/POPG (A) and POPG (D). Basic, polar, acidic and hydrophobic residues are coloured in blue, green, red and white respectively. The top view snapshot shows the tendency to aggregate for the eight peptides in POPE/POPG (B) or POPG (E). Heatmaps of average Cα-Cα distances for peptides in POPE/POPG (C) or POPG (F) show the stronger interactions that mediate assembly of aggregates.

Temporin L also adopts oligomeric configurations in both bilayers (Fig. 5). In POPE/POPG, again populations of monomer, dimer and trimer aggregates are observed (Fig. 5B) with self-association mediated by contacts between hydrophobic residues in the centre of the primary sequence (Phe5, Phe8, Leu9). Again, this was reproduced in both duplicates. In POPG, while again both monomers and dimers are observed, a higher order aggregate was detected in one of the duplicated simulations. This comprised four (Fig. 5D) or sometimes even five peptides (Fig. 5E) with a series of aligned, hydrophobic, phenylalanine residues (Phe1, Phe5, Phe8) found to be mediating the self-association (Fig. 5F). As can be seen from the snapshot, within this aggregate, peptides are oriented with the α-helix long axis oriented both tilted towards and perpendicular to the bilayer surface. In the duplicate simulation monomers and one trimer were observed, with self-association in that aggregate mediated by all residues between Phe8 and the C-terminus (FLGRIL).

### Aurein 2.5 and temporin L induce channel like activity in model membranes

As in our previous study^9^, we made use of the Port-a-patch® automated patch-clamp system from Nanion Technologies (Munich, Germany) to determine whether the two peptides could be distinguished on the basis of their disruption of either DPhPE:DPhPG (60:40 mol:mol) or DPhPG bilayers, mimicking Gram-negative and Gram-positive bacteria, respectively. For the temporin B analogues the technique determined substantial variation in the concentration of peptide required to induce conductance, the latency – the time taken for conductance to commence after addition of peptide, the duration of activity and whether the membrane was broken through the action of the AMP. In no cases was any characteristic channel like activity – well-defined events with discrete opening levels - observed. In contrast, in the present study both aurein 2.5 and temporin L were found to be capable of inducing channel like activity (Fig. 6). For aurein 2.5 this was observed in both membrane types (Fig. 6A/B) whereas for temporin L this was only found when challenging membranes formed from DPhPG (Fig. 6D).

**Figure 6.**
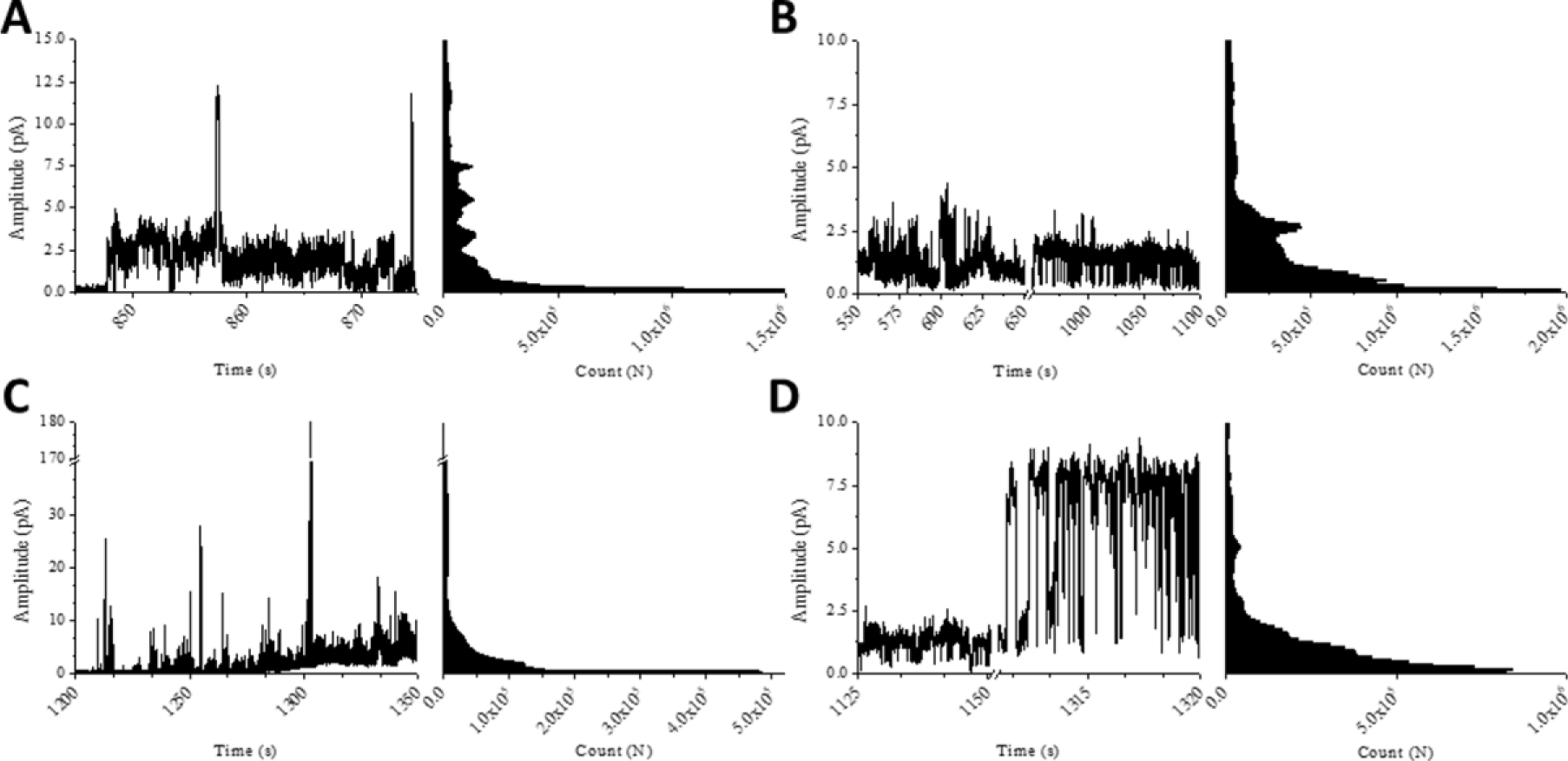
Membrane activity of the temporin B peptides. Representative current traces and all-points histograms for aurein 2.5 (A/B) and temporin L (C/D) in DPhPE/DPhPG (A/C) or DPhPG (B/D) membranes. Experiments were acquired with a holding potential of +50 mV and in the presence of 250 mM KCl, 50 mM MgCl_2_, 10 mM HEPES, pH 7.0. y-axis scaled accordingly to the different amplitudes for the different peptides.

Similar concentrations of aurein 2.5 and temporin L were required to induce channel activity in both membranes (Table 1) and this parameter therefore does not explain any of the observed differences in antibacterial potency. Membrane activity was observed to commence a little earlier for aurein 2.5 in both membranes (*p* < 0.05) but then the activity continued for both peptides for some minutes before the membrane ruptured (Fig. S4). For aurein 2.5 the measurements in both DPhPE/DPhPG and DPhPG membranes adopted a similar pattern with very many low amplitude events and few higher amplitude events (Fig. S5). Opening levels varied within and between replicate traces indicating that pores that form are both transient and irregular. Nevertheless, a set of discrete levels can be defined based on the frequency of certain amplitude events and this enables a measure of the conductance – a function of dwell time – which in turn can be used to estimate the pore size (Table 3). Three discrete levels were consistently identified for aurein 2.5 in both membranes with a slightly higher amplitude and conductance found in DPhPG membranes. The pore size estimate suggests pores approximately equal to the size of a chloride ion are formed by the peptide.

**Table 3.** Summery of channel-like activity detected at various opening levels. Pore size is estimated according to the formula provided in the materials and methods section.

| Peptide | Parameter |  |  |  |
| --- | --- | --- | --- | --- |
|  |  | Level 1 | Level 2 | Level 3 |
| Aurein 2.5<br>(DPhPE/DPhPG) | Amplitude<br>(pA) | $0.58 \pm 0.03$ | $1.29 \pm 0.07$ | $3.21 \pm 0.17$ |
| Aurein 2.5<br>(DPhPG) | | $0.83 \pm 0.03$ | $2.46 \pm 0.06$ | $5.43 \pm 0.04$ |
| Temporin L<br>(DPhPG) | | $0.89 \pm 0.02$ | **$25.38 \pm 0.14$** | - |
| Aurein 2.5<br>(DPhPE/DPhPG) | Conductance<br>(pS) | $11.54 \pm 0.52$ | $25.89 \pm 1.36$ | $64.24 \pm 3.38$ |
| Aurein 2.5<br>(DPhPG) | | $16.8 \pm 0.67$ | $49.22 \pm 1.30$ | $108.56 \pm 0.76$ |
| Temporin L<br>(DPhPG) | | $17.89 \pm 0.39$ | **$507.54 \pm 2.81$** | - |
| Aurein 2.5<br>(DPhPE/DPhPG) | Estimated pore<br>radius (nm) | $0.06 \pm 0.01$ | $0.09 \pm 0.02$ | $0.14 \pm 0.03$ |
| Aurein 2.5<br>(DPhPG) | | $0.07 \pm 0.01$ | $0.13 \pm 0.02$ | $0.19 \pm 0.01$ |
| Temporin L<br>(DPhPG) | | $0.08 \pm 0.01$ | **$0.43 \pm 0.03$** | - |

Similar to aurein 2.5, temporin L has channel like activity when challenging DPhPG membranes. For this peptide only two levels could be consistently detected but the amplitude and conductance associated with the higher of the two levels were substantially greater than that found and calculated for aurein 2.5 in the same membrane (Table 3). The resulting estimated pore size indicates that temporin L is capable of creating pores that are at least twice the diameter of a chloride ion. When challenging DPhPE/DPhPG, temporin L induces numerous high amplitude conductance events (Fig. 6C). However, these events are not well-defined or discrete. Instead, bursts of current (fast events seen as spikes) with a wide distribution of amplitudes are detected. This is similar to the activity observed for temporin B and its analogues,^9^ albeit with many events of much greater amplitude.

## Discussion

### Time-resolved techniques distinguish mechanistic features between the frog antimicrobial peptides aurein 2.5 and temporin L

Here we investigated whether two short, hydrophobic antimicrobial peptides, both derived from frog dorsal secretions and either known or predicted to adopt highly ordered α-helix conformations can be distinguished in their interactions with model membranes to better understand differences in their antimicrobial potencies and consequently the extent to which strategies for overcoming bacterial challenge may have converged in two distantly related frog species. Both aurein 2.5 and temporin L were confirmed as adopting highly ordered α-helix conformations in SDS, by CD and NMR, but also in more realistic membrane environments by CD. Both peptides were also capable of inducing channel-like conductance in model membranes and this should be contrasted with the membrane activity of temporin B,^9^ a similarly short and hydrophobic AMP that nevertheless adopts much less ordered α-helix conformation, which is incapable of inducing channel-like activity in the exact same conditions used for the present study. More detailed analysis of the channel conductance measurements and comparison of the results obtained from MD simulations however allow key differences in behaviour of the peptides to be revealed and suggest, not only that the two peptides operate with distinct mechanisms against the same membrane targets, but also that temporin L may adopt differing strategies for disrupting the plasma membranes of Gram-negative and Gram-positive bacteria. These two time-resolved techniques therefore appear useful in defining features of these AMPs that determine their potency. It is notable that the timescale of between 10 and 20 minutes for the appearance and duration of channel conductance is consistent with that observed for the attack of AMPs on bacteria reported elsewhere.^46,47^ It is currently not possible to sample such timescales using all-atom molecular dynamics simulations and the simulations here describe only the initial binding of the two peptides to the model bilayers. Nevertheless, key differences in the conformational flexibility of the two AMPs are shown. In the case of temporin L, the importance of the conformational flexibility found in the C-terminus region and the role of key residues in the formation of higher order aggregates is supported by a substantial body of previous work. We therefore consider the extent to which differences in antimicrobial and membrane activities can be related to the apparent function of individual elements of the primary sequence whose behaviour has been partly revealed here at the molecular level.

### The high relative potency of temporin L against Gram-positive bacteria is associated with larger pores and higher conductance due to the formation of higher order aggregates

Interpretation of the findings from the MD simulations is substantially aided by previous structure-activity relationship and biophysical studies, most notably of temporin L.^17–20,22–24^ In particular, substitutions for Gln3, Gly10 and various replacements of the three phenylalanine residues have been made and their effect on antibacterial potency, cell cytotoxicity, haemolysis and anti-endotoxin activity determined.

The C-terminal segment, Gly10-Leu13, and the N-terminus to Gln3 are shown here to experience substantial conformational flexibility. The C-terminal segment penetrates less into the hydrophobic core of model membranes than the remainder of the peptide while Gln3 is less buried when binding to POPE/POPG bilayers. The C-terminal segment and Gln3 were the most exposed to solvent in SDS micelles in a previous NMR study^19^ hence there is agreement that temporin L is located at the interfacial region of the membrane with these sections of the sequence having specific roles. Further support for this comes from the effect of substituting Gly10 for either leucine or proline.^17^ Replacement of Gly10 with leucine substantially increases the propensity to form α-helix and leads to abrogation of the antibacterial activity against both Gram-positive and Gram-negative species and drastically increased haemolytic potential. In contrast, replacement of Gly10 with proline produced a peptide with negligible haemolytic potential but at the costs of a significant reduction in antibacterial activity. The conclusion that helical conformation at the C-terminus is essential for antibacterial activity^17^ can be refined to state that conformational flexibility in the C-terminal region is essential for antibacterial activity and this is afforded by glycine and not proline. As seen here for aurein 2.5 and previously^10^ for magainin 2, glycine alone is insufficient to induce conformational flexibility in an AMP sequence and the residues located close to Gly10 can be expected to contribute.

Gln3 has also been replaced with proline,^17,19^ which has little effect on the antimicrobial activity but reduces a little the haemolytic potential, and lysine,^22,24^ which increases the antiendotoxin capability of the peptide and produced a modest improvement in antibacterial activity. In these analogues the first two, hydrophobic residues were not altered, and the retention of activity is consistent with the essential role, shown here, for these residues in the initial binding as these penetrate the most into the hydrophobic core. Increasing the flexibility around the third residue may alter binding to membranes of differing composition while increasing the cationicity may enhance binding to the anionic surface of bacterial plasma membranes without compromising insertion; as was observed for analogues of temporin B.^9^

The importance of an identified phenylalanine zipper motif^23^ can also now be considered in the context of its role, shown here, in mediating contacts between self-associated peptides in dimers and higher order aggregates. The hydrophobic residues, in particular the three phenylalanines are involved in the initial binding to the membrane and the stabilisation of peptide aggregates in the membrane. Phe5 and Phe8 have been substituted for either alanines or leucines.^23^ Substitution of either residue with alanine substantially reduces antibacterial activity, cytotoxicity and haemolytic potential with substitution of Phe5 having greater impact. Substituting leucine for either Phe5 or Phe8 has little impact on antibacterial activity or cytotoxicity but reduces a little the haemolytic potential. Interestingly however, substitution of leucine for both Phe5 and Phe8 leads to total loss of antibacterial activity against *S. aureus* 25923 while activity against *E. coli* 25922 and *P. aeruginosa* BAA 427 is retained; haemolysis and cytotoxicity are increased. This indicates that both Phe5 and Phe8 are required for activity against Gram-positive *S. aureus* and that, combined, they have a specific role when temporin L challenges this, and related, species which cannot be substituted by other hydrophobic residues. In the present study, higher order aggregates and pores with diameters greater than twice the size of a chloride ion were observed only in membranes designed to model the Gram-positive plasma membrane i.e. uniformly negatively charged. Further, in membranes designed to model the Gram-negative plasma membrane, neither higher order aggregates nor regular channel like activity was observed. We therefore hypothesise that in Gram-positive plasma membranes only, higher order temporin L aggregates can form and these lead to relatively larger channels and this underpins high antibacterial potency against such species. By extension, temporin L uses a different mechanism of permeabilising the plasma membrane of Gram-negative bacteria which does not require the formation of higher order aggregates or regular channels, but which nevertheless induces very high amplitude conductance.

### Aurein 2.5 forms low order aggregates and acts via low conductance channel like permeabilisation of both Gram-negative and Gram-positive plasma membranes

In comparison with temporin L, less is known about the role of individual residues in aurein 2.5. Further, although both peptides are confirmed as preferring ordered α-helix conformations, aurein 2.5 behaves differently to temporin L in models of both Gram-positive and Gram-negative membranes. Notably however, glycine, that confers conformational flexibility to the C-terminal segment of temporin L does not have the same effect in aurein 2.5 while hydrophobic residues, which include two phenylalanines but also three valines and two leucines, mediate the formation of dimers or trimers but not higher order aggregates. Channels that are formed are small and induce relatively low conductance.

There are two glycines in the primary sequence of aurein 2.5, Gly14 and Gly11, with the latter located in a position where it might be expected to interrupt a long section of α-helix. In contrast with temporin L, where conformational flexibility is greater but confined to segments located at the N-or C-terminus, conformational flexibility is more modest and is evenly distributed along the length of the peptide. Our recent comparison of conformational flexibility in pleurocidin and magainin 2 found substantial difference in conformational flexibility between the two peptides despite very similar positioning of glycine residues^10^ and, taking both studies together, we can conclude that this important property will be dependent both on positioning of the glycines themselves but also on contributions from residues in their vicinity.

The hydrophobic phenylalanines that are distributed in a heptad zipper motif and are responsible for mediating peptide self-association in temporin L are, in aurein 2.5, located close to either the N-or C-terminus. Instead, hydrophobic valine residues, Val 6, Val9 and Val10, are distributed throughout the aurein 2.5 primary sequence and perform the role of mediating self-association. Whether or not the geometry of aurein 2.5 allows for the formation of higher order aggregates, the selection of valine over phenylalanine in these positions may also preclude their formation. The selection of an ability to form low or higher order aggregates may well impact on the ability to cause high conductance membrane permeabilisation but it may impact other parameters. Notably aurein 2.5 induces channel like activity much more rapidly than temporin L in DPhPG membranes and a little less peptide is needed. Future work may explore whether indeed higher order aggregates are needed for high conductance channel like activity and whether this is associated with a slower onset of channel activity.

## Conclusion

Two AMPs from distantly related frogs were found to adopt seemingly identical, ordered α-helix conformations in model membranes. MD simulations and conductance measurements indicated however that there are fundamental differences between the two AMPs in how they bind to and disrupt these membranes. One AMP, temporin L, was also found to differ fundamentally in how it permeabilises models of membranes of either Gram-negative and Gram-positive bacteria and that these differing strategies were dependent on key contributions from individual amino acid residues in the primary sequence.

## ASSOCIATED CONTENT

**Supporting Information**. Secondary structure analysis of the peptides as determined by far-UV CD in POPE/POPG or POPG lipid small unilamellar vesicles, further analysis of the MD simulations and representations of the key parameters from the Port-a-patch® data are provided as supplementary figures S1-S5.

## AUTHOR INFORMATION

### Author Contributions

GM, DAP and AJM designed the study. GM, PMF, and AJM wrote the main manuscript text and prepared all figures. PMF and CDL performed and/or analysed atomistic simulation data. SSB provided materials. GM, HA and RAA performed structural studies. GM, VBG, HA, TTB and AFD performed and/or analysed CD measurements. GM and VBG performed and analysed patch-clamp measurements. GM, CH, MC and MJS conducted antimicrobial susceptibility testing. All authors approved the manuscript.

### Notes

The authors declare no competing interests.

#### Acknowledgment

NMR experiments described in this paper were produced using the facilities of the Centre for Biomolecular Spectroscopy, King’s College London, acquired with a Multi-user Equipment Grant from the Wellcome Trust and an Infrastructure Grant from the British Heart Foundation. CDL acknowledges the stimulating research environment provided by the EPSRC Centre for Doctoral Training in Cross-Disciplinary Approaches to Non-Equilibrium Systems (CANES, EP/L015854/1). PMF is supported by a Health Schools Studentship funded by the EPSRC.

## REFERENCES

1 Andersson, D. I., Hughes, D. and Kubicek-Sutherland, J. Z. (2016) Mechanisms and consequences of bacterial resistance to antimicrobial peptides. Drug Resist. Update. 26, 43–57

2 Zasloff, M. (2002) Antimicrobial peptides of multicellular organisms. Nature. 415, 389–395

3 Giuliani, A., Pirri, G., Bozzi, A., Di Giulio, A., Aschi, M. and Rinaldi, A.C. (2008) Antimicrobial peptides: natural templates for synthetic membrane-active compounds. Cell. Mol. Life Sci. 65, 2450–2460

4 Hancock, R.E., and Diamond, G. (2000) The role of cationic antimicrobial peptides in innate host defences. Trends Microbiol. 8, 402–410

5 Haney, E.F., and Hancock, R.E.W. (2013) Peptide design for antimicrobial and immunomodulatory applications. Biopolymers. 100, 572–583

6 Jenssen, H., Hamill, P., and Hancock, R.E.W. (2006) Peptide antimicrobial agents. Clin. Microbiol. Rev. 19, 491–511

7 Nguyen, L.T., Haney, E.F., and Vogel, H.J. (2011) The expanding scope of antimicrobial peptide structures and their modes of action. Trends Biotechnol. 29, 464–472

8 Simmaco, M., Mignogna, G., Canofeni, S. Miele, R., Mangoni, M.L. and Barra, D. (1996) Temporins, antimicrobial peptides from the European red frog *Rana temporaria*. Eur. J. Biochem. 242, 788–792

9 Manzo, G, Ferguson, P.M., Gustilo, V.B., Ali, H., Bui, T.T., Drake, A.F., Atkinson, R.A., Batoni, G., Lorenz, C.D. Phoenix, D.A. and Mason, A.J. (2018) Minor sequence modifications in temporin B cause drastic changes in antibacterial potency and selectivity by fundamentally altering membrane activity. bioRxiv DOI: 10.1101/312215

10 Amos S-B.T.A., Vermeer, L.S., Ferguson, P.M., Kozlowska, J., Davy, M., Bui, T.T., Drake, A.F., Lorenz, C.D. and Mason, A.J. (2016) Antimicrobial peptide potency is facilitated by greater conformational flexibility when binding to Gram-negative bacterial inner membranes. Sci. Rep. 6, 37639–37651

11 Kozlowska, J., Vermeer, L.S., Rogers, G.B., Rehnnuma, N., Amos S-B.T.A., Koller, G., McArthur, M., Bruce, K.D. and Mason, A.J. (2014) Combined systems approaches reveal highly plastic responses to antimicrobial peptide challenge in *Escherichia coli*. PLoS Pathog 10 (5), e1004104

12 Mangoni, M.L., Di Grazia, A., Cappiello, F., Casciaro, B., and Luca, V. (2016) Naturally occurring peptides from *Rana temporaria*: Antimicrobial properties and more. Curr. Top. Med. Chem. 16, 54–64.

13 Rinaldi, A.C., Mangoni, M.L., Rufo, A., Luzi, C., Barra, D., Zhao, H., Kinnunen, P.K.J., Bozzi, A., Di Giulio, A. and Simmaco, M. (2002) Temporin L: antimicrobial, haemolytic and cytotoxic activities, and effects on membrane permeabilization in lipid vesicles. Biochem. J. 368, 91–100

14 Rosenfeld, Y., Barra, M., Simmaco, M., Shai, Y., and Mangoni, M.L. (2006). A synergism between temporins towards Gram-negative bacteria overcomes resistance imposed by the lipopolysaccharide protective layer. J. Biol. Chem. 281, 28565–28574

15 D’Abramo, M., Rinaldi, A.C., Bozzi, A., Amadei, A., Mignogna, G., Di Nola, A. and Aschi, M. (2006) Conformational behaviour of temporin A and temporin L in aqueous solution: a computational/experimental study. Biopolymers 81, 215–224

16 Mangoni, M.L., Papo, N., Barra, D., Simmaco, M., Bozzi, A., Di Giulio, A. and Rinaldi, A.C. (2004) Effects of the antimicrobial peptide temporin L on cell morphology, membrane permeability and viability of *Escherichia coli*. Biochem. J. 380, 859–865

17 Mangoni, M.L., Carotenuto, A., Auriemma, L., Saviello, M.R., Campiglia, P., Gomez-Monterrey, I., Malfi, S., Marcellini, L., Barra, D., Novellino, E. and Grieco, P. (2011) Structure-activity relationship, conformational and biological studies of temporin L analogues. J. Med. Chem. 54, 1298–1307

18 Merlino, F., Carotenuto, A., Casciaro, B., Martora, F., Loffredo, M.R., Di Grazia, A., Yousif, A.M., Brancaccio, D., Palomba, L., Novellino, E., Galdiero, M., Iovene, M.R., Mangoni, M.L. and Grieco, P. (2017) Glycine-replaced derivatives of [Pro^3^,DLeu^9^]TL, a temporin L analogue: Evaluation of antimicrobial, cytotoxic and hemolytic activities. Eur. J. Med. Chem. 139, 750–761

19 Carotenuto, A., Malfi, S., Saviello, M.R., Campiglia, P., Gomez-Monterrey, I., Mangoni, M.L., Gaddi, L.M., Novellino, E. and Grieco, P. (2008) A different molecular mechanism underlying antimicrobial and hemolytic actions of temporins A and L. J. Med. Chem. 51, 2354–2362.

20 Saviello, M.R., Malfi, S., Campiglia, P., Cavalli, A., Grieco, P., Novellino, E. and Carotenuto, A. (2010) New insight into the mechanism of action of the temporin antimicrobial peptides. Biochemistry 49, 1477–1485

21 Giacometti, A., Cirioni, O., Ghiselli, R., Mocchegiani, F., Orlando, F., Silvestri, C., Bozzi, A., Di Giulio, A., Luzi, C., Mangoni, M.L., Barra, D., Saba, V., Scalise, G. and Rinaldi A.C. (2006) Interaction of antimicrobial peptide temporin L with lipopolysaccharide *in vitro* and in experimental rat models of septic shock caused by Gram-negative bacteria. Antimicrob. Agents Chemother. 50, 2478–2486.

22 Srivastava, S. and Ghosh J.K. (2013) Introduction of a lysine residue promotes aggregation of temporin L in lipopolysaccharides and augmentation of its antiendotoxin property. Antimicrob. Agents Chemother. 57, 2457–2466

23 Srivastava, S., Kumar, A., Tripathi, A.K., Tandon, A. and Ghosh, J.K. (2016) Modulation of anti-endotoxin property of temporin L by minor amino acid substitution in identified phenylalanine zipper sequence. Biochem. J. 473, 4045–4062

24 Farrotti A, Conflitti P, Srivastava S, Ghosh JK, Palleschi A, Stella L, Bocchinfuso G. (2017) Molecular dynamics simulations of the host defense peptide temporin L and its Q3K derivative: an atomic level view from aggregation in water to bilayer perturbation. Molecules 22, E1235

25 Rozek, T., Wegener, K.L., Bowie, J.H., Olver, I.N., Carver, J.A., Wallace, J.C. and Tyler, M.J. (2000) The antibiotic and anticancer active aurein peptides from the Australian Bell Frogs *Litoria aurea* and *Litoria raniformis*. Eur. J. Biochem. 267, 5330–5341

26 Dennison, S.R., Morton, L.H.G., Shorrocks, A.J., Harris, F.H. and Phoenix, D.A. (2009) A study on the interactions of Aurein 2.5 with bacterial membranes. Colloids Surf. B Biointerfaces. 68, 225–230

27 Dennison, S.R., Morton, L.H.G. and Phoenix, D.A. (2012) Role of molecular architecture on the relative efficacy of aurein 2.5 and modelin 5. Biochim. Biophys. Acta. 1818, 2094–2102

28 Dennison, S.R., Harris, F., Morton, L.H. and Phoenix, D.A. (2013) Antimicrobial activity of aurein 2.5 against yeasts. FEMS Microbiol. Lett. 346, 140–145

29 Dennison, S.R., Morton, L.H., Harris, F. and Phoenix, D.A. (2014) The interaction of aurein 2.5 with fungal membranes. Eur. Biophys. J. 43, 255–264

30 Wang, J., Mura, M., Zhou, Y., Pinna, M., Zvelindovsky, A V., Dennison, S.R. and Phoenix, D.A. (2014) The cooperative behaviour of antimicrobial peptides in model membranes. Biochim. Biophys. Acta. 1838, 2870–2881

31 Zhang, P., Liang, D., Mao, R-L., Hillis, D.M., Wake, D.B. and Cannatella, D.C. (2013) Efficient sequencing of Anuran mtDNAs and a mitogenomic exploration of the phylogeny and evolution of frogs. Mol. Biol. Evol. 30, 1899–1915.

32 Keller, R. (2004) The Computer Aided Resonance Assignment Tutorial. Zurich: Cantina Verlag

33 Dynamo software: The NMR Molecular Dynamics and Analysis System. Available online at: http://spin.niddk.nih.gov/NMRPipe/dynamo

34 Guerry, P., and Herrmann, T. (2012) Comprehensive Automation for NMR Structure Determination of Proteins. Methods in Molecular Biology. 831, 429–451

35 Schwieters, C., Kuszewski, J., Tjandra, N. and Clore, G.M. (2003) The Xplor-NIH NMR molecular structure determination package. Journal of Magnetic Resonance. 160 (1), 65–73

36 Schwieters, C., Kuszewski, J. and Clore G.M. (2006) Using Xplor-NIH for NMR molecular structure determination. Prog. Nucl. Magn. Reson. Spectrosc. 48, 47–62

37 Manzo, G., Serra, I., Pira, A., Pintus, M., Ceccarelli, M., Casu, M., Rinaldi, A.C. and Scorciapino, M.A. (2016) The singular behavior of a β-type semi-synthetic two branched polypeptide: three-dimensional structure and mode of action. Phys. Chem. Chem. Phys. 18, 30998–31011

38 Abraham, M. J., Murtola, T., Schulz, R., Páll, S., Smith, J. C., Hess, B. and Lindahl, E. (2015) GROMACS: High performance molecular simulations through multi-level parallelism from laptops to supercomputers. SoftwareX. 1, 19–25

39 Best, R.B., Zhu, X., Shim, J., Lopes, P.E.M., Mittal, J., Feig, M. and MacKerell, Jr, A.D. (2012) Optimization of the additive CHARMM all-atom protein force field targeting improved sampling of the backbone φ, ψ and side-chain χ1 and χ2 dihedral angles. J. Chem. Theory Comput. 8 (9), 3257–3273

40 Huang, J., and MacKerell, A.D. (2013) CHARMM36 all-atom additive protein force field: validation based on comparison to NMR data. J. Comput. Chem. 34, 2135–2145

41 Lee, J., Cheng, X., Swails, J.M., Yeom, M.S., Eastman, P.K., Lemkul, J.A., Wei, S., Buckner, J., Jeong, J.C., Qi, Y., Jo, S., Pande, V.S., Case, D.A., Brooks III, C.L., MacKerell Jr., A.D., Klauda, J.B. and Im, W. (2016) CHARMM-GUI input generator for NAMD, GROMACS, AMBER, OpenMM, and CHARMM/OpenMM simulations using the CHARMM36 additive force field. J. Chem. Theory Comput. 12 (1), 405–413

42 Angelova, M. and Dimitrov, D.S. (1986) Liposome electroformation. Faraday Discuss. Chem. Soc. 81, 303–311

43 Angelova, M. and Dimitrov, D.S. (1988) A mechanism of liposome electroformation. Trends in Colloid and Interface Science II, ed Degiorgio V (Springer, Berlin), 59–67

44 Angelova, M. (2000) Giant vesicles. Perspectives in Supramolecular Chemistry, eds Luisi PL, Walde P (Wiley-Interscience, Chichester, UK), 1st Ed, 27–36

45 Tosatto, L., Andrighetti, A.O., Plotegher, N., Antonini, V., Tessari, I., Ricci, L., Bubacco, L. and Dalla Serra, M. (2012) Alpha-synuclein pore forming activity upon membrane association. Biochim. Biophys. Acta 1818, 2876–2883.

46 Sochacki, K.A., Barns, K.J., Bucki, R. and Weisshaar, J.C. (2011) Real-time attack on single *Escherichia coli* cells by the human antimicrobial peptide LL-37. Proc. Natl. Acad. Sci. U S A. 108, E77–E81

47 Choi, H., Yang, Z. and Weisshaar, J.C. (2017) Oxidative stress induced in *E. coli* by the human antimicrobial peptide LL-37. PLoS Pathog. 13, e1006481.

